# Oomycete communities in lowland tropical forest soils vary in abundance and are composed of saprophytes and pathogens of seeds and seedlings of multiple plant species

**DOI:** 10.1101/2024.02.25.580666

**Authors:** K.D. Broders, H.D. Capador-Barreto, G. Iriarte, S. J. Wright, H. Espinosa, M. Baur, M.A. Lemus-Peralta, E. Rojas, E.R. Spear

## Abstract

**Premise:** The soils in lowland tropics are teeming with microbial life which can impact plant community structure and diversity through plant-soil feedbacks. While bacteria and fungi have been the focus of most studies in the tropics, the oomycetes may have an outsized effect on seed and seedling health and survival, given their affinity for environments with increased precipitation and temperature.

**Methods:** We assessed the diversity and pathogenicity of oomycete species present in a lowland tropical forest in Panama. We used both a culture dependent leaf-baiting assay and culture independent soil DNA metabarcoding methods to quantify zoospore abundance and species diversity. A subset of the isolates from the baiting assay were used to evaluate pathogenicity and aggressiveness on seedlings of three tree species.

**Key results:** Oomycetes are ubiquitous and common members of the soil microbial community in lowland tropical forests and zoospore abundance was far greater compared to similar studies from temperate and mediterranean forests. We also observed variation in oomycete species ability to infect host plants. Species of *Pythium* were more aggressive, while species of *Phytopythium* caused less disease but were more diverse and commonly isolated from the soil. Finally, we found that individual hosts accumulate a distinct oomycete community and was the only factor that had an effect community structure.

**Conclusions:** Collectively, these finding demonstrate that oomycetes are ubiquitous, host-generalist pathogens and saprophytes, that have the potential to impact seed and seedling survival in lowland tropical forests

## INTRODUCTION

The microbial communities in lowland tropical forests are known to have an impact on plant community structure (Mangan et al. 2010, Bagchi et al. 2014, Delavaux et al. 2023). The bacteria and fungi have generally been the focus of these interactions between the soil microbial community and plant species. However, the oomycetes, which are fungal-like organisms, are increasingly being observed in culture independent studies in tropical ecosystem (Mahe et al. 2017, Oliverio et al. 2020, Delavaux et al. 2023). While oomycetes resemble fungi in their morphology and growth as filamentous threads known as mycelia and nutrient acquisition through absorption, they are classified with the stramenopiles, which includes the brown algae and diatoms that have lost plastids and are phylogenetically distant from kingdom Fungi (Beakes et al. 2012, Keeling and Burki 2019). Oomycetes occur as part of healthy ecosystems, are an important component to soil biodiversity and play important roles in ecosystem processes (Gomez-Aparico 2012, Bever 2015, Geisen 2015) by functioning as saprotrophs that degrade organic material, pathogens and parasites of invertebrates, vertebrates, plants and fungi (Beakes et al. 2012, Jiang et al. 2013, Spies et al. 2016, Grover and Barkoulas 2021). As pathogens, these organisms may limit host population sizes thereby affecting ecosystems processes, including primary and secondary production, biogeochemical cycles, disturbance regimes, and physical structure (Garnas et al. 2011, Cobb et al. 2012, Sato et al. 2012, Avila et al. 2016).

The most well-studied genera include *Phytophthora* and *Pythium* which contain many plant pathogens with the potential to infect and damage a wide range of hosts. While much of our knowledge of these genera comes from agricultural systems, recent surveys of oomycetes in natural forests have unveiled a number of novel species (Jung et al. 2020, 2018), including several pathogens with the capacity to infect and impact growth of trees at all growth stages, often resulting in mortality. These oomycetes grow as mycelia in the soil and produce motile spores that move through water in and on the surface of soil. Because oomycetes can readily move through the soil and infect plants, they are excellent seed, seedling, and root pathogens and are believed to drive patterns of conspecific negative density dependence (CNDD) in temperate and Mediterranean tree species (Packer and Clay 2000, 2004, Dominguez-Begines et al. 2021), and may contribute to the maintenance of species in lowland tropical forests as soil-borne pathogens (Mangan et al. 2010). While many of the pathogenic oomycetes associated with agricultural crops and temperate tree species have been identified and their lifecycles, epidemiology, reproduction cycles and even genomes elucidated, very little is known about the diversity and ecological roles of oomycetes in tropical forests.

Lowland tropics are an ideal environment for the growth, reproduction, and dispersal of oomycetes. Many species of *Pythium, Phytopythium* and *Phytophthora* survive as oospores or chlamydospores in soil and dead plant tissue. These survival structures can then germinate to either grow as mycelia or produce sporangia. The sporangia can then be dispersed by wind or rain, directly infect a host upon contact, or produce zoospores with two flagella that can swim in free water (Agrios 2005). These zoospores are chemotactically and electrotactically attracted to the surfaces of host plants (Khew and Zentmyer 1973, Latijnhouwers et al. 2004). Zoospores swim until reaching a host, at which point they shed their flagella, encyst, and firmly attach themselves to the plant’s surface via secretion of adhesion molecules (Deacon and Donaldson 1993).

Oomycetes have been found to infect seeds and seedlings causing both pre- and post-emergent damping-off of a variety of plant species and have been implicated in causing density- and distance-dependent mortality in the lowland tropical forests of Panama (Augspurger and Kelly 1984, Augspurger and Wilkinson 2007, Mangan et al. 2010, Sarmiento et al. 2017). Damping off is often amplified at high seed and seedling densities because diseased individuals enhance both the growth and the dispersal of the pathogen via mycelia and zoospores. This results in an exponential rate of spore production, dispersal, and infection of nearby seeds and seedlings until seeds or seedlings are at densities sufficiently low or spaced at distances sufficiently large to limit dispersal (Burdon and Chilvers 1975, Augspurger and Kelly 1984) or grow and accrue resources to resist infection through ontogenic resistance (Simon et al. 2014, Ampt et al. 2022).

Initial studies evaluating the oomycetes causing disease on young seedlings in the lowland tropics have only occurred within the last 25 years (Davidson 2000, Augspurger and Wilkinson 2007). While studying the frequency-dependent selection of pathogens on seedlings of *Anacardium excelsum*, *Tetragastris panamensis* and *Callophylum longifolium,* Davidson (2000) was able to isolate and prove pathogenicity of several oomycetes including *Phytophthora* (*Phytoph.*) *heveae* and an assemblage of *Pythium* species, including *Pythium splendens*, *Phytopythium* (*Phytopy.*) *chamaehypon and Phytopy. vexans*. Davidson (2000) also studied the effect of *Phytoph. heveae* and *Phytopy. vexans* on density-dependent disease dynamics among seedlings of *A. excelsum*. The pathogen species responded differently to host density and seasonality. *Phytoph. heveae* was associated with high seedling density and mortality sites, whereas *Phytopy. vexans* was not and infections by *Phytopy. vexans* were observed throughout the wet season. This demonstrated that oomycete species vary in their aggressiveness, or rate of infection and host mortality, as well as their ability to disperse and spread between plants. This also suggests that the species of oomycetes inhabiting a location impact the strength of observed CNDD. This observed variation in aggressiveness and mortality supports existing evidence that oomycete pathogens can be bio-, hemibio- and necrotrophs (Fawke et al. 2015). *Phytophthora* species tend to be hemibiotrophic, initially growing as a biotroph (deriving energy from living cells) but later switch to a nectrotroph (from dead or dying cells) as the host begins to die. The genera *Pythium* and *Phytopythium* tend to be necrotrophic and often live saprophytically (decomposing dead matter) in the soil and in plant litter. *Pythium* and *Phytopythium* species are most frequently associated with diseased seeds, seedlings and young roots that have not developed physical resistance barriers to pathogen infection.

A global analysis of soil protists found that oomycetes were the second most abundant parasitic protist and the most abundant plant parasitic protist (Oliverio et al. 2020). Parasitic protists, including oomycetes, were especially abundant in tropical soils, up to 38% of the community (Oliverio et al. 2020). There was also a strong positive correlation between oomycete abundance and moisture and temperature, with abundance peaking in tropical sites. In combination, Oliverio et al. (2020) and Mahe et al (2017) suggest parasitic oomycetes have an important role in tropical ecosystems as saprophytes and parasites of seeds, seedlings, and roots of young trees. This is supported by previous experiments on seedling mortality (Augspurger and Kelly 1984, Davidson 2000, Augspurger and Wilkinson 2007) and seeds of *Trema micrantha* and *Apeiba membranaeae* infected with oomycetes in a lowland tropical forest (Sarmiento et al. 2017). While these studies demonstrate the potential importance of oomycete species in driving patterns of CNDD through seed and seedling mortality, we possess limited information on the species diversity, pathogenicity and aggressiveness, and host preference of oomycetes.

The aim of this study was to understand the species diversity, distribution, abundance and pathogenicity of soil-borne oomycetes in a lowland tropical forest in Panama. We used traditional culture-based baiting (briefly, immersing soil in sterile water and allowing zoospores to swim up and infect floating leaf pieces) and culture-independent, DNA-based metabarcoding to assess oomycete communities from soil samples collected from 22 sites across the Barro Colorado Nature Monument representing six soil types and ten tree species. The goals of this project were to: 1) compare and contrast the efficacy of these methods for estimating oomycete diversity; 2) assess the pathogenicity and host range of the recovered oomycetes; and 3) determine the factors structuring oomycete communities in the soil of a lowland tropical forest.

## MATERIAL AND METHODS

### Study site

We characterized soil-borne oomycete communities within Barro Colorado Nature Monument (BCNM) Republic of Panama (9.063°N; 79.50°W) (Figure S1). BCNM is a 5600-hectare reserve that includes Barro Colorado Island (BCI) and five surrounding peninsulas. The semideciduous forests are classified as tropical moist forests, with ca. 2600 mm rainfall annually and a mean temperature of 27°C (Windsor 1990). The soils of BCI are mapped and diverse deriving from four parent materials of distinct geological formations and six major soil forms (Yavitt 2024).

### Sample collection

Soil samples were collected from 22 sites in BCNM, including 7 sites on BCI and 8 sites on Gigante Peninsula in May and June of 2019, respectively (Fig S1). Sampling on BCI focused on collecting samples from different soil types and soil samples were collected near recent soil profiles representing 5 types (Suppl. Table 1; (Yavitt 2024)). The soil sample was also taken at least 1.5 m from the nearest adult tree, and the identity of the nearest adult tree recorded. There were numerous root pieces in the soil sample, which likely represented more tree species than the nearest individual recorded in Table 1. Sample sites on Gigante Peninsula were selected for proximity to adults of five tree species. On Gigante, soil was collected beneath two individuals each of the following species: *Simarouba amara*, *Heisteria concina*, *Hirtella triandra*, and *Dailium guianense.* An additional 7 sites on BCI were collected from beneath *Virola nobilis* (syn. = *V. surinamensis*) trees (December 24 and 25, 2019), specifically 3 male and 4 female trees (Suppl. Table 1; Fig. S2A) used in a previous study that found potential genotype-specific effects of pathogens on plant species coexistence (Eck et al. 2019). Soil was collected from 3 evenly spaced locations per tree. Each location was 1.5 m from the trunk, and we avoided sampling next to other adult trees and/or saplings. At each sampling location, we used a bleach-sterilized hand spade to disturb and homogenize the soil in a small depression (15 cm diameter, 20 cm deep). All soil was temporarily (<48 hours) stored in an opaque bin at 4°C. A subsample was collected and stored at -80°C for future DNA extraction. The remaining soil was stored in an opaque bin at 4°C prior to baiting.

**Table 1.** Analysis of variance outputs for percent of bait-leaves infected.

|  | DF | Df.res. | F-value | p-value |
| --- | --- | --- | --- | --- |
| Intercept | 1 | 161.877 | 20.5228 | 1.14E-05 |
| rep | 4 | 151.001 | 0.8152 | 0.5173 |
| species | 10 | 17.412 | 0.316 | 0.9661 |
| rep*species | 28 | 151.001 | 1.0023 | 0.4703 |

### Culture-based isolation and quantification of oomycetes from soil

Leaf baiting has been demonstrated to be effective for isolating and quantifying motile zoospores produced by oomycetes from soil and water samples (Rollins et al. 2016, Masikane et al. 2019). Briefly, soil is immersed in sterile water and oomycete zoospores swim up from the soil and infect floating leaf pieces via chemotaxis and because zoospores are negatively geotropic (Eden et al. 2000). This method has been used to isolate oomycetes from a variety of substrates (soil, roots, water), is effective at recovering a range of species, and has the advantage of selecting for those oomycetes that respond to the presence of and infect plant tissue, selecting for putative plant pathogens rather than saprotrophs and vertebrate and invertebrate pathogens (Balci et al. 2010, Jung et al. 2017, Jung et al. 2020, Sarker et al. 2023). In our experiments, we placed 50 grams of soil per sample in an opaque container with 250 ml of autoclave-sterilized distilled water (Fig. S2B). Fifteen surface sterilized (2 min, 70% EtOH; 3 min, 10% bleach) baits (leaf pieces approximately 1 cm^2^) were floated on the water surface abaxial side down. We used expanding (immature) mango (*Magnifera indica)* leaves. Preliminary experiments using leaves from mango, *Theobroma cacao*, *Anacardium excelsum*, and *Ochroma pyramidale* trees showed no significant difference in the number of oomycete infections across the four tree species. All containers were covered to ensure dark conditions (Fig. S2C) and stored at room temperature (∼24°C). After 72 hours, leaf baits were collected, surface sterilized, and placed on the selective growth medium potato dextrose agar (PDA) with the antibiotic rifampicin (+rif) and the antifungal pimaricin (+pim) (Fig. S2D and E). Plates were stored in the dark and checked for oomycete growth from infected leaf tissue after 48 and 96 hours. Unique morphotypes emerging from infected tissue pieces were hyphal-tip transferred to new PDA plates to obtain pure cultures (Fig. S2F). For *V. nobilis*, the three soil samples collected per tree were processed individually with 15 leaf pieces per soil sample, resulting in 45 leaf pieces per tree. For the soil samples collected near other tree species, the single bulk soil sample was divided into four containers with 10 leaf pieces per, resulting in 40 leaf pieces per collection site. For each soil sample, we considered the percent of bait leaves for which at least one oomycete isolate emerged to be a proxy for zoospore incidence. We tested whether zoospore incidence varied across tree species and sites using Linear Mixed-Effects Model (lmer) in R v. 4.2.2.

### Oomycete identification

Since we recovered over 600 oomycete isolates by baiting, we selected a subset of isolates that represented all unique morphological groups baited from each soil. This resulted in a set of 154 isolates identified using DNA sequence data. For DNA extraction, 20-70 mg of fresh oomycete mycelia was collected with a plastic tip and put into OPS Diagnostic prefilled bead tubes (100-& 400-micron silica beads) with extraction buffer (7.45 % (w/v) KCl, 0.1 M Tris HCl (pH 8.0), 0.01M EDTA (pH 8.0). Tubes were incubated for 15 min at 65°C and then the mycelial mass was pulverized for 20 seconds with a mini bead beater. The cell lysate was centrifuged at 10,000 rpm for 10 minutes, and the supernatant was decanted directly into tubes containing 0.3ml (300μl) of Isopronanol and mixed by inverting the tube 50 times. Thereafter, the tube was centrifuged at 12,000 rpm for 10 minutes, the supernatant discarded, and the pellet washed with 70% ethanol and centrifuged at 12000 rpm for 5 minutes. After the ethanol evaporated, pellets were dissolved in MilliQ water.

The internal transcribed spacer (ITS) region of the ribosomal DNA and the cytochrome oxidase I (COI) region of mitochondrial DNA have both proven to be effective markers for species level identification of oomycetes (Robideau et al. 2011). The ITS region was amplified and sequenced using the ITS5 and ITS4 primer pair (White et al. 1990). The COI fragment was amplified and sequenced using the oomycete-specific primers OomCoxILevup and Fm85mod to amplify 727 bp from the COI mitochondrial DNA (Robideau et al. 2011). In some cases, an alternative reverse primer, OomCoxI-Levlo was used with OomCoxI-Levup, amplifying a slightly smaller 680 bp fragment of COI, that still completely overlaps with the standard DNA barcode used in other groups (Robideau et al. 2011). PCR products were visualized by gel electrophoresis using 2% agarose gel stained with GelRed. All amplicons were purified and sequenced in both directions using the amplification primers at the MMUL, Smithsonian Tropical Research Institute (STRI; Panama City, PA) or at the Sequencing facility at the National Center for Agricultural Utilization Research (Peoria, IL, USA). Sequences and chromatograms were quality checked and forward and reverse sequence reads aligned and consensus sequence made using Sequencher v. 5.2.4 (Gene Codes Corporation). Pronounced double peaks were considered as heterozygous positions and labelled according to the IUPAC (International Union of Pure and Applied Chemistry (https://iupac.org) coding system. All sequences generated in this study were deposited in GenBank and accession numbers are given in Supplemental Table 2.

### Phylogenetic analysis

For phylogenetic analyses, the sequences obtained in this study were complemented with publicly available sequences of vouchered isolates from *Phytophthora*, *Phytopythium* and *Pythium* sourced from the GenBank Nucleotide Collection. The ITS and COI sequences used in the analyses were aligned using the MAFFT v.7 (Katoh and Standley 2013) plugin within the Geneious Prime® software by the G-INS-I algorithm. The ITS and COI alignments were also concatenated and aligned to determine the phylogenetic structure of oomycetes recovered during this study. The Maximum Likelihood (ML) analyses were completed with IQ-Tree 1.6.12 (Nguyen et al. 2015)(http://www.iqtree.org/) and Bayesian inference completed with MrBayes v.3.2.7 (Ronquist et al. 2012) plug-in in Geneious Prime. For Maximum Likelihood (ML) analyses the ITS, COI and combined datasets were aligned with MAFFT v. 7 (Katoh and Standley 2013). ModelFinder (Kalyaanamoorthy et al. 2017) was used to identify the best-fit model of molecular evolution for each partition based on the Bayesian information criterion scores (Chernomor et al. 2016). The phylogeny was reconstructed with IQ-Tree v. 1.6.12 using 2,000 standard nonparametric bootstrap replicates. For Bayesian inference, we used the general time reversible (GTR) model selected for the entire unpartitioned alignment, with the likelihood parameters setting (lset) number of substitution types (nst) = 6, with a proportion of sites invariable and the rest drawn from the gamma distribution (rate=invgamma). Four independent analyses, each starting from a random tree, were run under the same conditions for the combined gene alignment. Three hot and one cold chains of Markov Chain Monte Carlo iterations were performed. Analyses were run with 1,000,000 generations with sampling every 100 generations. The first 250,000 generations were discarded as the chains were converging (burnin period). Resulting trees were visualized with MEGA 11 v. 11.0.11 (Tamura et al. 2021).

#### Illumina MiSeq amplicon sequencing

In addition, to the culture-dependent method, we used metabarcoding and oomycete-specific primers to amplify and sequence the oomycete communities in each soil sample. DNA was extracted using DNeasy Powersoil Pro kit (Qiagen, Germantown MD). DNA quality and quantity was assessed using a Nanodrop. Oomycete communities were amplified using a two-stage PCR protocol. Locus-specific primers used for PCR1 included the Illumina sequencing primer sequence on the 5’ ends. We amplified region 1 of the internal transcribed spacer (ITS) barcode using oomycete-specific primers ITS1oo and ITS2 as described previously (Riit et al. 2016). We used Platinum 2X Mastermix (Thermo) in PCR reactions with a final volume of 12.5µl. The reaction was amplified as follows: initial denaturation at 95°C for 3 min, 34 cycles of denaturation at 95°C for 30s, annealing at 55°C for 30s, extension at 72°C for 1 min, followed by final extension at 72°C for 10 min. PCR2 used 2ul of PCR1 as template and added on remaining Illumina adaptors and index sequences. PCR2 products were cleaned and normalized using PCR Normalization plates (CharmBiotech, USA) and pooled libraries concentrated using AMPure beads (Beckman Coulter, USA). Libraries were sequenced on an Illumina MiSeq with 250bp paired end reads at the MMUL facility at STRI (Panama City, Panama).

#### Processing microbial community data

Reads were trimmed of forward and reverse primers using cutadapt (v1.18) (Martin 2011) following an initial filtering step that removed reads with ambiguous bases. Primer sequences with more than 12% error rate (–error-rate = 0.12) were discarded. Reads were then processed using DADA2 (Callahan et al. 2016) (v1.16.0) within R (v4.1.0). Reads were dropped from the data set if they had three or more expected errors (maxEE = 2), at least one base with very low quality (truncQ = 2), or at least one position with an unspecified nucleotide (maxN = 0). Reads were dereplicated before inferring amplicon sequence variants (ASVs). Paired-end reads were merged and read pairs that did not match exactly across at least 12 base pairs (minOverlap = 12) were discarded. We retained amplicons between 100 and 450 base pairs. Reads were then screened for chimeras (method = consensus). For taxonomic classification, we used the naïve Bayesian classifier (Wang et al. 2007) against the general release of the UNITE database for eukaryotes from July 2023 (Abarenkov et al. 2023). To reduce computational time, we filtered the database to include only organisms in clade Stramenopila, where all oomycetes belong.

Subsequent data analysis was performed in R using the package *phyloseq* (McMurdie and Holmes 2013). Two samples, originating from two *V. nobilis* trees had extremely low diversity values likely due to laboratory errors and excluded from subsequent analysis. We filtered ASVs with less than 10 reads in total to explore the distribution of the putative pathogens in the soil. To explore the abundance of the isolated oomycetes we determined the presence and abundance of ASVs across the 10 *V. nobilis* adults and heterospecific trees in surrounding forests (BCI and the adjacent Gigante peninsula). The ASV sequences from the metabarcoding study were blasted against the ITS sequences from isolated oomycetes using BLASTn (Camacho et al. 2009) using a 99% match identity and 50% coverage to ensure the accuracy of the matches.

### Oomycete pathogenicity

We selected nineteen oomycete isolates to experimentally evaluate pathogenicity. These isolates were recovered from the baiting assay, specifically soils collected adjacent to 9 tree species, and included four isolates of *Pythium splendens* as well as nine isolates from the *Phytopythium vexans* complex. These experiments were initiated prior to sampling near *V. nobilis* trees and STRI facilities were closed during the height of the Covid-19 pandemic, so we could not evaluate any *Phytophthora* isolates. We could not test the oomycete isolates against the tree species present at the soil source because seeds were unavailable. Thus, we evaluated pathogenicity with seedlings of *Ochroma pyramidale*, *Swietenia macrophylla,* and *Theobroma cacao*. All three tree species are present in BCNM, but again were not near any sampling location. Consequently, our pathogenicity assays evaluated: (1) whether the isolates are generalist pathogens and (2) the potential host range. Seeds were manually cleaned, and surface sterilized by submersion in 10% commercial sodium hypochlorite (3 min), followed by 70% ethanol (3 min) and 3 rinses with sterile water. Seeds were sown in germination trays with individual compartments containing a mix of 2:1 soil:sand, which was autoclaved twice for 45 minutes. Recently germinated seedlings at the first cotyledon stage were inoculated.

From January-March 2020, we evaluated the pathogenicity of cultured oomycetes using a rice grain inoculum protocol used previously to assess fungal pathogenicity on tropical tree seedlings (Spear and Broders 2021). Inoculum consisted of autoclave sterilized rice grains uncolonized (control) or colonized by a single oomycete. For the later, sterile rice grains were aseptically placed on actively growing cultures of each oomycete isolate. Each recently germinated seedling was transplanted to an individual cone-tainer (5cm diameter x 20cm depth) and filled with steam-sterilized soil. Prior to transplanting, four uncolonized rice grains (control) or four grains colonized by one of the 19 oomycete isolates tested were placed in a depression in each cone-tainers soil and the seedling was planted with its roots in contact with the rice grains. For each treatment, we inoculated 10 *O. pyramidale* seedlings, 10 *S. macrophylla* seedlings, and 5 *T. cacao* seedlings. Five weeks after inoculation, seedlings were harvested, above ground symptoms scored on an ordinal rating scale (1 = healthy, 2=mild wilting and chlorosis, 3=moderate wilting and stunted growth, 4=Severe wilting and stunted growth, and 5 = Dead) and the presence/absence of lesions on roots and stems noted (Figure S3). Root lesions were plated on the selective medium PDA +rif +pim with the hope of reisolating the oomycete inoculated onto the seedling thereby satisfying Koch’s postulates.

## RESULTS

### Prevalence of Oomycetes from soil

Oomycetes were recovered from all 22 soil sites sampled in this study using both culture-dependent leaf baiting and culture-independent metabarcoding methods. Considering all sites, we obtained 642 oomycete isolates from leaf baiting (Suppl. Table 1). However, zoospore incidence (measured as percent baits infected) varied widely across the 22 sites ranging from 4-93% of baits infected, with 12 sites having greater than 50% of baits infected (Fig. 1). There was no effect of tree species on zoospore incidence, but there was a significant effect (P < 0.001) of site (Table 2). Based on previous estimates of zoospore abundance associated with leaf-bait infection incidence (Rollins et al. 2016, Masikane et al. 2019), the 8 sites in this study with 25-50% infection incidence had an estimated 100-1,000 zoospores/ml of water. The 12 sites with >50% infection incidence had an estimated 1,000 – 100,000 zoospores/ml of water.

**Figure 1.**
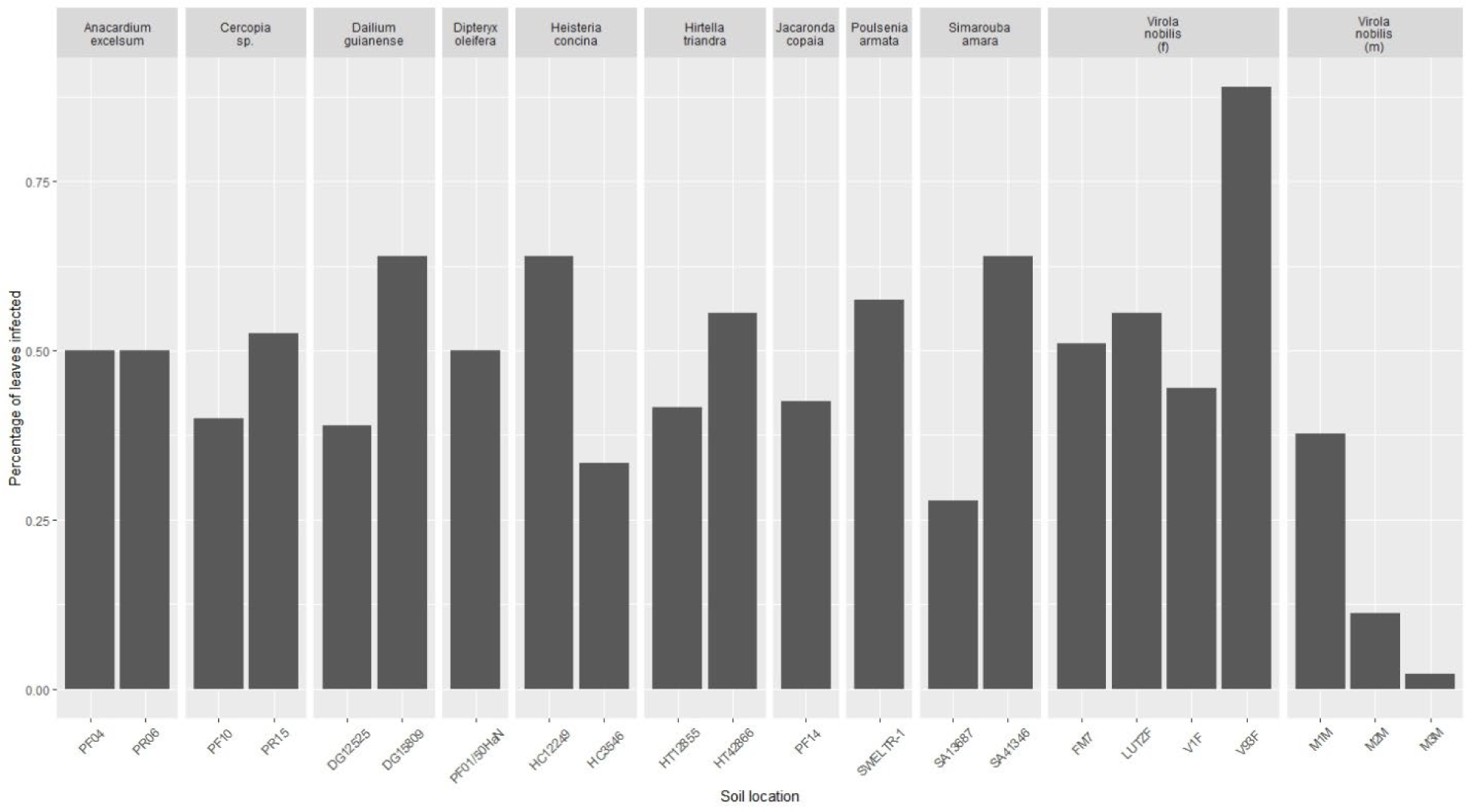
Zoospore abundance, measured as the percent of bait leaves infected from 22 soils near 11 host species collected in a lowland tropical forest in Panama.

### Oomycete diversity and distribution

Twenty-four unique oomycete species were identified in the soils of the sampled sites using baiting. Isolates of the genus *Phytopythium* were the most abundant and diverse across all sites, except soils near *V. nobilis* adults, and included the detection of 14 putative new species (Figure 2, Figure S4). Isolates in the genus *Phytophthora*, including *Phytoph. heveae, castanaea,* and *panamensis*, were the most frequently recovered oomycetes near *V. nobilis* adults, but they were not recovered from any sites without *V. nobilis* (Figure 3). No oomycete species were found at all 22 sites. The most frequently recovered species were *Phytopythium vexans* complex sp. 1 (PVC 1) and PVC 6, both of which were isolated from 9 and 8 sites, respectively (Figure 3). *Phytophthora castanae* and *Pythium splendens* were the most frequently isolated species not in the *Phytopy. vexans* complex and were isolated from 4 and 5 sites, respectively. However, *Phytoph. castanaea* was only recovered from near *V. nobilis* adult trees, whereas *Py. splendens* was recovered from near 5 tree species (Figure 3).

**Figure 2.**
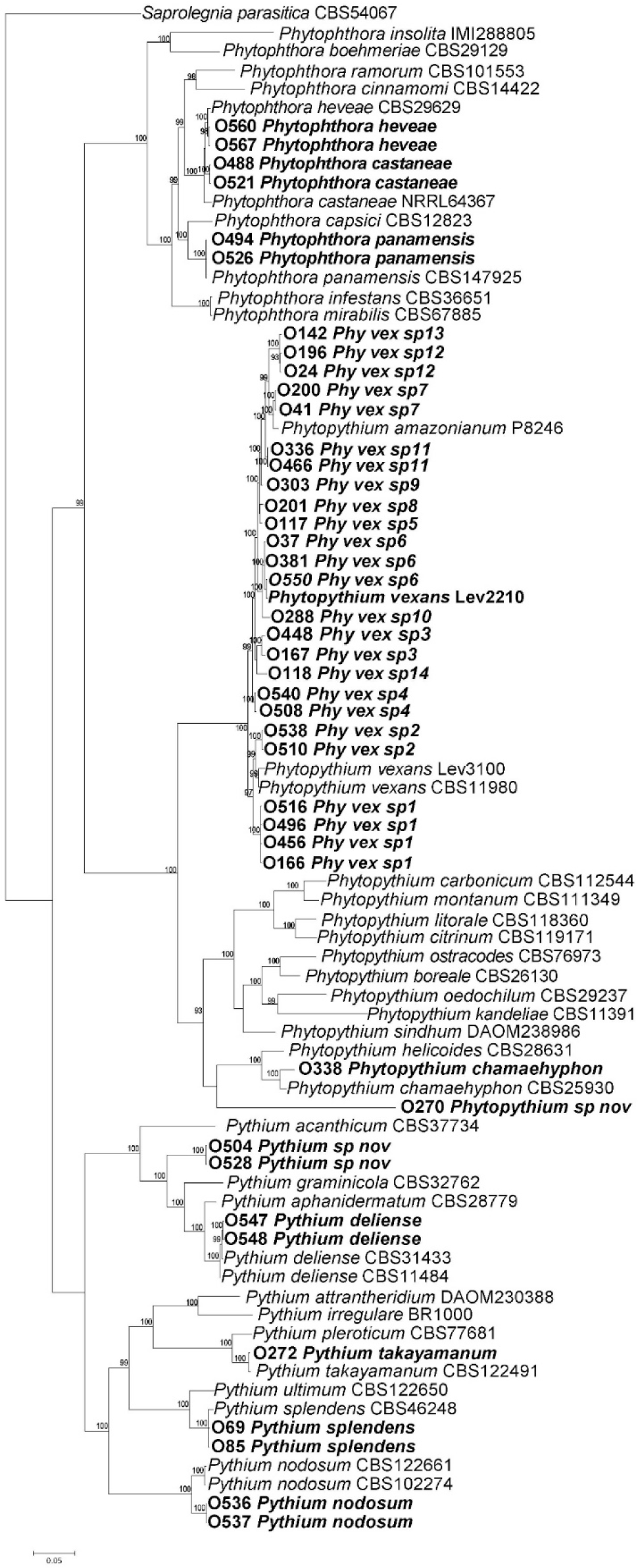
Phylogenetic diversity of oomycetes species baited from soil in a lowland tropical forest in Panama derived from a Bayesian phylogenetic analysis of a concatenated data set of ITS and cox1 gene sequences of isolates from this study and representative species from other genera of the Peronosporaceae and Pythiaceae. Species in bold were isolated during this project.

**Figure 3.**
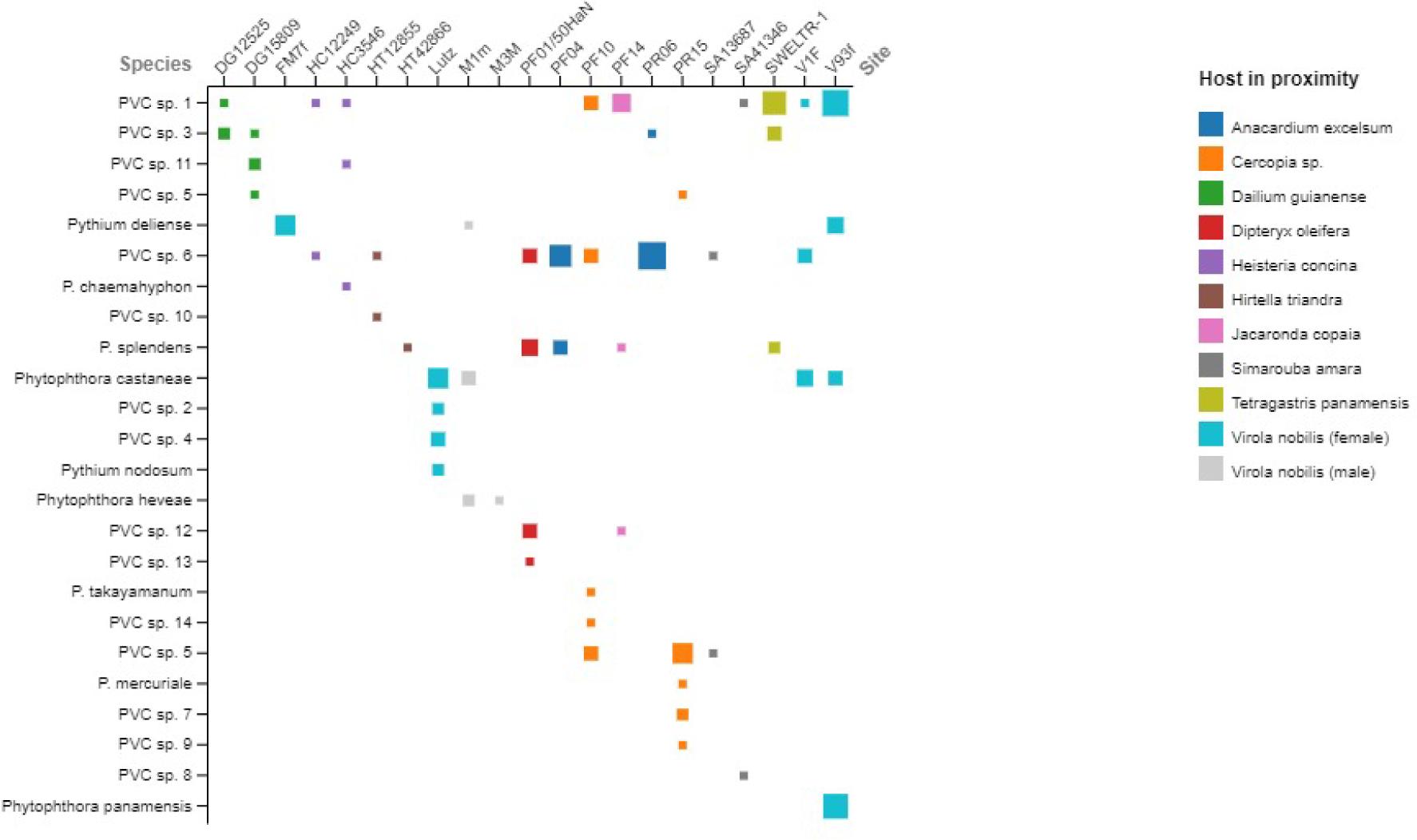
Abundance and distribution of oomycetes species in a lowland tropical forest in Panama. Oomycetes were baited from the top 15 cm of soil and the closest host species recorded. The size of the square is proportional to the abundance of a species at a single site.

The metabarcoding data broadly supported the baiting results. Greater diversity was detected with metabarcoding, identifying 499 oomycete ASVs. Members of the genus *Phytopythium* (including *Oomycota gen. Incertae sedis*, *Peronosporales gen. incertae sedis,* see note on taxonomy below) were found to be the most abundant plant-associated genus at 17 of the sites, *Pythium/Globisporangium* (*Globisporangium* is considered part of *Pythium* sensu latu and includes *P. splendens*) was the most abundant at 4 sites and one site had similar relative abundances of these two groups (Figure 4). Unlike the baiting assay, the metabarcoding was able to detect the presence of *Phytophthora* at 10 sites not associated with *V. nobilis* trees. (Figure 4). There was no effect of nearest tree species on oomycete community, but oomycete communities varied significantly across individual trees based on the metabarcoding data. We could not assess community composition with the baiting results due to the limited number of isolates identified to species for each soil sample. However, we observed significant variation in zoospore infection incidence across sampling sites (Table 2), supporting our metabarcoding-based conclusion that oomycete communities vary across sites.

**Figure 4.**
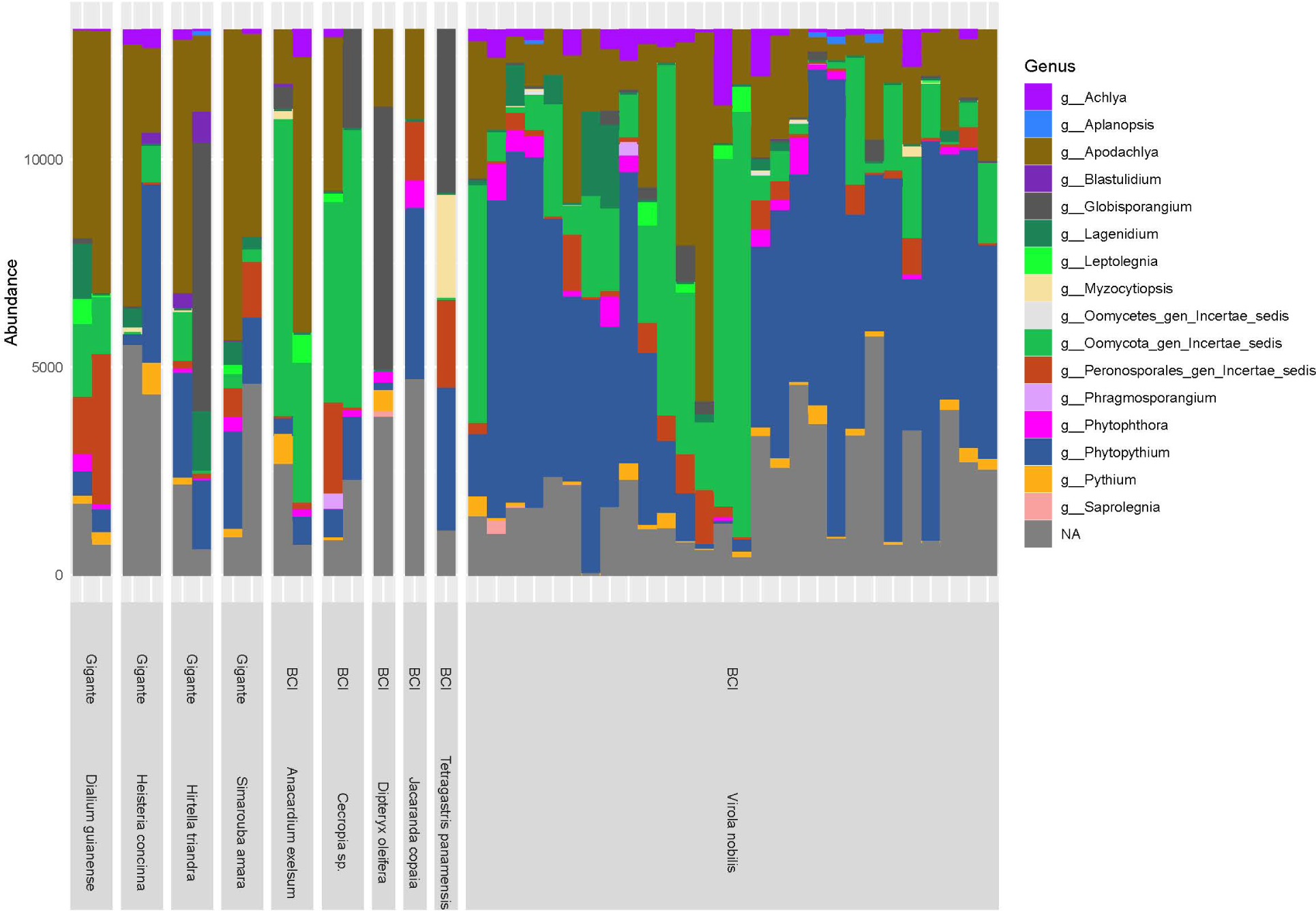
Oomycete community composition from soil collected near 11 tree species in a lowland tropical forest in Panama. Samples are grouped by location (BCI, Gigante) and host species. Relative abundance of Oomycete genera is denoted in different colors.

The UNITE database was able to assign taxonomy to genus for most oomycete ASVs. However, there were several highly abundant ASVs classified as *Oomycota gen. Incertae sedis* and *Peronosporales gen. incertae sedis* at several sites and we wanted to determine if these were also found in the baiting assay. We found that UNITE incorrectly assigned several ASVs that were >99% similar to *Phytopythium* from the baiting study to other genera including *Oomycota gen. Incertae sedis* and *Peronosporales gen. incertae sedis* (Suppl Table 3). This is likely the result of many novel species of *Phytopythium* identified in this study, and the lack of representation of many members of the Oomycota in the UNITE database.

Previous studies on BCI and other regions of the tropics and subtropics have found multiple species within the *Phytopythium vexans* species complex (PVC) that cause disease on seedlings, infect seeds, and cause root rots (Figure 5). In a previous study by Sarmiento et al. (2017) at least 7 species within the PVC were recovered from infected seeds of *Trema micrantha* and *Apeiba membranaceae* on BCI (Figure 5). Of those 7 species, 5 were also recovered during this study. Many of the other PVC isolates with similarity to isolates from this study were recovered from diseased tree crop roots and stems such as Areca nut, Avocado, Durian, Papaya, Persimmon, Raspberry, as well as from native forests (Figure 5). There are also at least six distinct clades within the PVC that have only been recovered from forests in Panama (Figure 5).

**Figure 5.**
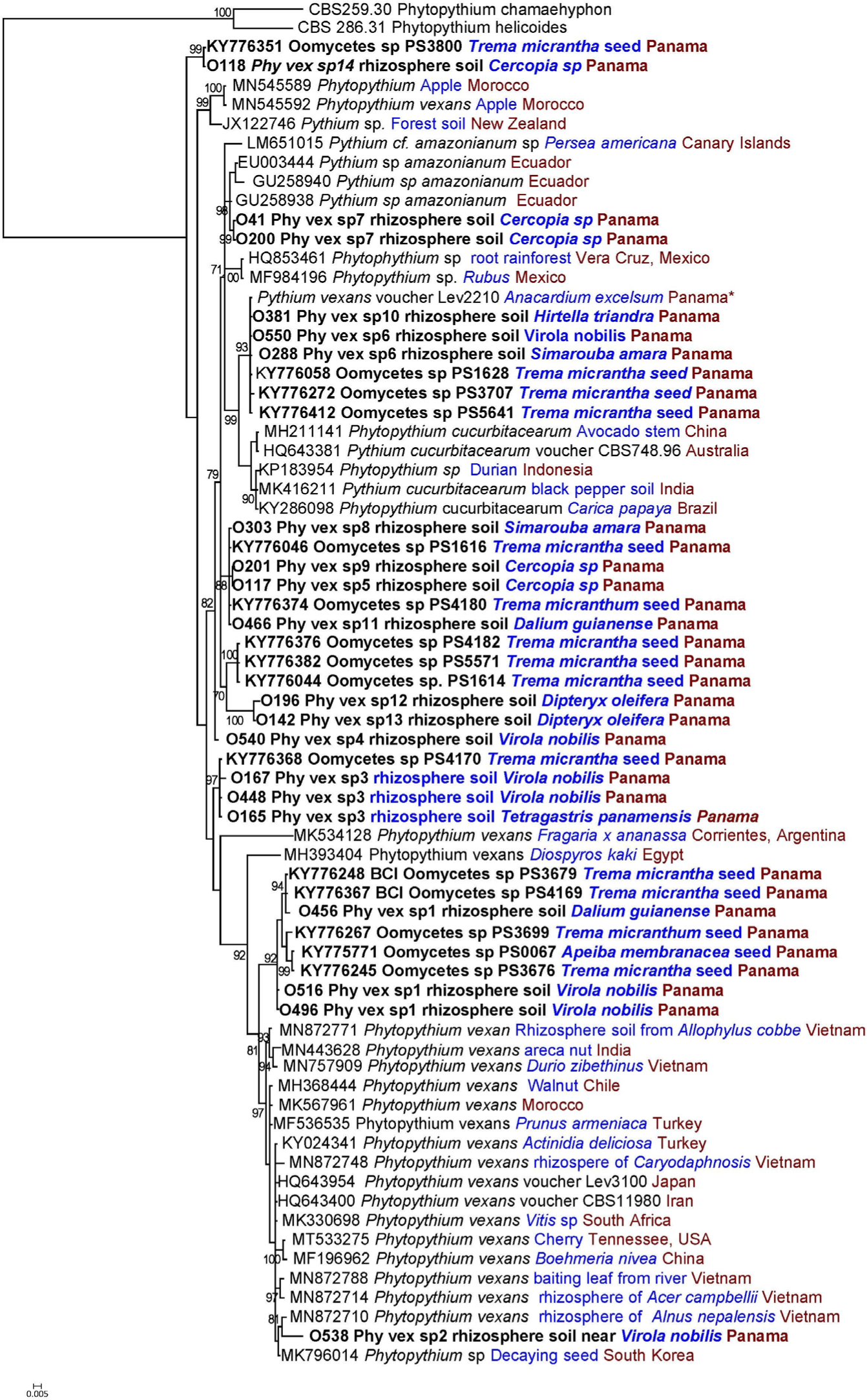
Phylogenetic diversity of *Phytopythium vexans* species complex strains associated with the soil, seeds and roots derived from Bayesian analysis of the ITS region of rDNA from sequences generated from this project, additional sequences from Barro Colorado Island, Panama generated in Sarmiento et al. 2017 (PS numbers) and sequences imported from Genbank that were >97% similar to at least one of the *Phytopythium* sequences generated in this study. Isolates collected on BCI during this study and Sarmiento et al. 2019 are denoted in bold. Information on host/substrate is indicated in blue text and geographic origin in red text. Strain CBS 119.80 is the living ex-type of the species *Phytopythium vexans*.

### Oomycete pathogenicity

*Pythium splendens* was the most aggressive oomycete tested and exhibited a broad host range. It caused severe seedling wilting and mortality for all three tree species on which it was inoculated (Figure 6; Fig. S3). All *Phytopythium* isolates tested were weakly pathogenic or not pathogenic on seedlings of the three tropical tree species. The *Phytopythium* isolates evaluated rarely caused mortality during the 5-week pathogenicity assays, but they did routinely cause lesions on roots and the lower stem, near the soil line, of seedlings of multiple tree species. We could not assess the long-term impacts of the root and stem infections on survival and growth.

**Figure 6.**
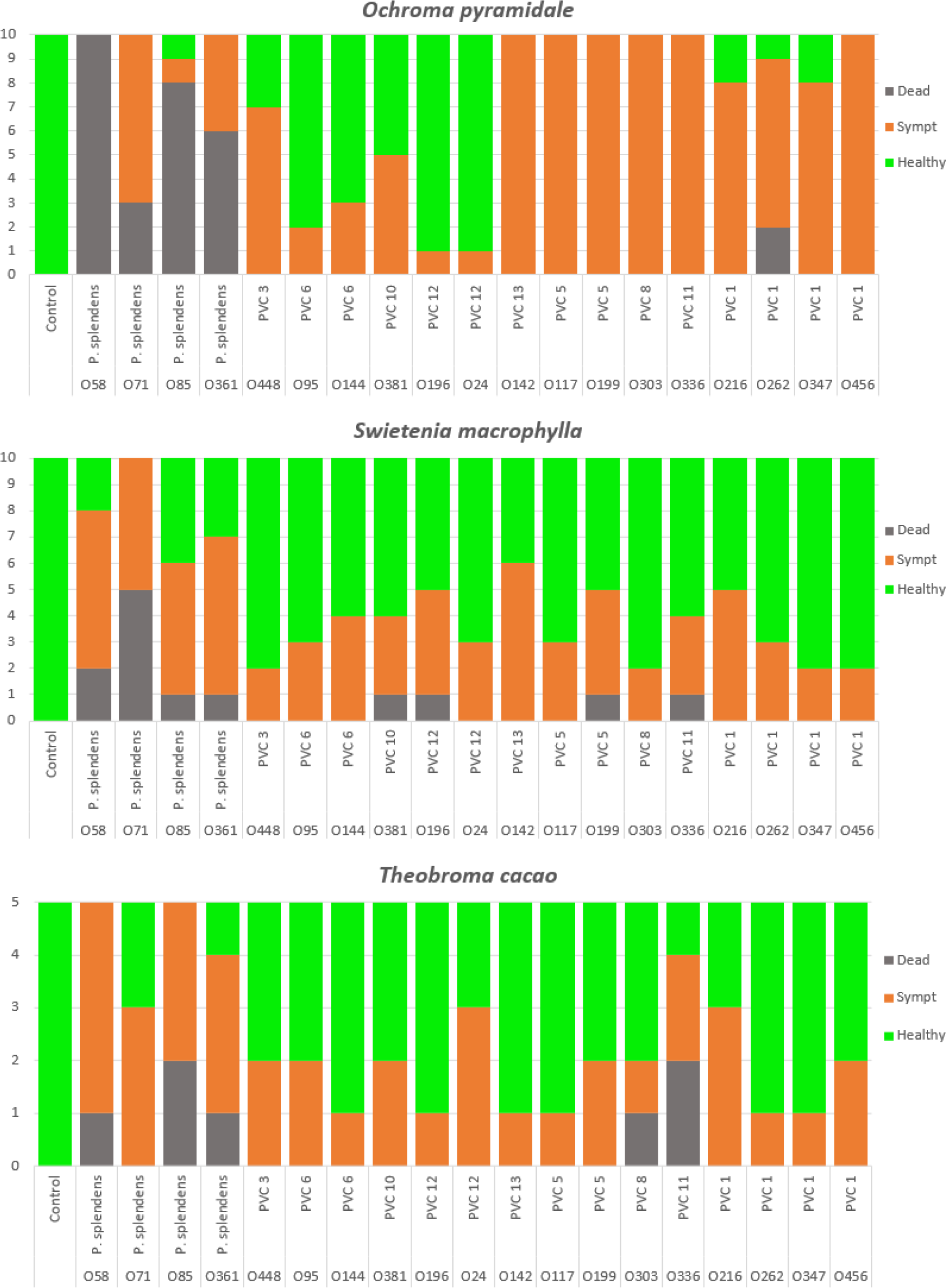
Pathogenicity of 19 oomycete strains recovered from soil in a lowland tropic forest in Panama. Ability to cause mortality (dark grey), lesions on the roots (orange), and no disease (green) were assessed on 10 individual seedlings of a) *Ochroma pyrimadale*, b) *Swietenia macrophylla*, and 5 individual seedling of c) *Theobroma cacao*.

## DISCUSSION

Our results highlight the advantages of utilizing both culture dependent and independent methods for studying oomycete diversity in the tropics. Metabarcoding provided a more precise characterization and assessment of species diversity and relative abundance at each site, while baiting facilitated the detection and characterization of multiple putative new species and quantification of infective propagules (zoospores). Both approaches uncovered low diversity of *Phytophthora* and *Pythium* compared to temperate and highland tropical forests, as well as a greater than expected diversity of *Phytopythium* across the study sites. The zoospores from *Phytophthora* species were only recovered in the baiting assay from soil near *V. nobilis* trees sampled at the end of the rainy season in December, whereas *Pythium* and *Phytopythium* species were associated with multiple tree species and were found in equal numbers from sites sampled at the early part of the rainy-season (June) and end of the rainy season (December). However, the metabarcoding data detected the presence of *Phytophthora* species near 8 of the 10 tree species assessed in this study (Figure 4). This suggests that *Phytophthora* species may require more time to become active and increase zoospore production after rewetting of soil after a prolonged dry period. Also, as *Phytophthora* species tend to be hemibiotrophic, they may require a plant host to infect that will allow for production of secondary inoculum including mycelia and zoospores. This would occur throughout the rainy seasons as many seeds begin germinated in July through November resulting in peak zoospore production of *Phytophthora* species in December. Additional surveys will need to be conducted to confirm this pattern.

### Prevalence of Oomycetes in lowland tropical forest soil

The frequency at which oomycetes were isolated from the soils on BCI and Gigante was greater than has been observed in temperate forests. Oomycetes were present in all soil samples and >25% individual leaf baits were infected for 20 of the 22 sites sampled on BCI and Gigante. These results are similar to those from other lowland tropical forests in French Guyana where oomycetes were recovered from 92 of 93 sites (Legeay et al. 2020) and lowland and highland tropical and sub-tropical forests in Vietnam where oomycetes were recovered from 54 of 55 sites across the country (Jung et al. 2020). In contrast, in temperate forests in the central and eastern United States the incidence of bait-leaf infections ranged from 0-23% (Balci et al. 2007). In contrast with our results, only 21% of soils from mixed hardwood forests in Ohio, USA yielded isolates (Balci et al. 2007). Similar to the Ohio study, only 42% of the 137 soil samples tested from near *Quercus, Betula, Acer* and *Fagus* species in Pennsylvania, USA yielded oomycete isolates (Bily et al. 2022). Even when soils were sampled near trees with visible *Phytophthora* infections in a declining beech forest in Switzerland, only 70% of soils yielded infected bait leaves (Ruffner et al. 2019). This suggests that oomycetes are more abundant and active in tropical versus temperate forests. The primary driver of the stark contrast between tropical and temperate forests is likely the higher annual precipitation and temperatures in tropical forests, as oomycete relative abundance is positively correlated with both mean annual precipitation and mean annual temperature based on a global study of soil eukaryotic diversity which included both temperate and tropical forest sites (Oliverio et al. 2020).

Based on zoospore quantification results from a similar baiting assay (Rollins et al. 2016, Masikane et al. 2019), there is the potential for an estimated 100 – 100,000 zoospores per gram of water-saturated soil on BCI and Gigante. This represents substantial potential for infection of seeds and seedlings in times of high precipitation. Due to their biology and production of swimming spores, plant-pathogenic oomycetes have the ability to increase their dispersal, growth and infection of otherwise healthy hosts as moisture availability increases (Jung 2009, Sturrock et al. 2011). For this reason, oomycetes are commonly associated with significant disease epidemics during periods of increased precipitation (Thompson et al. 2013, Campoverde et al. 2017). The species assessed in this study are either able to cause disease ranging from minor root lesions to wilting and mortality on multiple tree species. This observed lack of host specificity and capacity to produce hundreds of thousands of infective propagules indicate that oomycetes are a major source of seed and seedling disease in periods of high precipitation in lowland tropical forests and elevated disease in wetter versus drier forests, as previously observed across a precipitation gradient in Panama (Spear et al. 2015). Increased oomycete and fungal disease potential positively correlated with precipitation may be an important driver of observed increases in plant diversity with soil moisture in the lowland tropics of Panama (Lebrija-Trejos et al. 2023). Lebrija-Trejos et al. (2023) determined the negative density-dependent interactions among conspecifics were stronger and increased diversity in wetter years, suggesting the moisture availability enhances diversity indirectly through moisture-sensitive, density-dependent conspecific interactions, including increased disease pressure from pathogens like the oomycetes that positively respond to increases in precipitation (Oliverio et al. 2020, Delavaux et al. 2021).

Oomycetes were ubiquitous across all soil types and near all tree species sampled. Isolates of *Phytopythium* were the most abundant and diverse across all sites and we detected 14 putative new species (Fig. S4). In contrast, the presence and relative abundance of isolates from the genera *Pythium* and *Phytophthora* varied across sites, but *Pythium* was more aggressive than *Phytopythium* in pathogenicity experiments. All *Phytopythium* isolates tested were weakly pathogenic or not pathogenic on the seedlings of three tropical tree species but may still negatively impact seed and seedling survival. While *Phytopythium* are generally less aggressive pathogens compared to *Phytophthora* and *Pythium*, many species are increasingly being recognized as important root rot pathogens that impact long-term health of various plant species (Wang et al. 2015, Fichtner et al. 2016, Noireung et al. 2020, Baysal-Gurel et al. 2021, Li et al. 2021). While these root and stem infections may not lead to seedling death in the short-term, they may affect the long-term fitness and survival of the seedling. Members of the *Phytopythium vexans* complex (PVC) were the most commonly isolated pathogen of diseased *Anacardium excelsum* and *Tetragastris panamensis* seedlings on BCI (Davidson et al. 2000) and multiple OTUs from the PVC were recovered from diseased seeds on BCI (Sarmiento et al. 2017). In fact, based on the sequence data, 5 of the 7 oomycete OTUs infecting seeds in Sarmiento et al. (2017) were also present in the soils used in this project and detected by leaf baiting (Figure 6).

The sporadic distribution and abundance of *Pythium* and *Phytophthora* indicate these species may accumulate under individual trees or in specific soil types, and in conjunction with their virulence, suggests they may contribute to patterns of conspecific negative density dependence. The aggressiveness, relatively broad host range and sporadic distribution of *Py. splendens* represents an example of a multihost pathogen with the potential to promote coexistence and enhance diversity if infection differentially affects each host (Hersh et al. 2012, Sedio and Ostling 2013) and if the abiotic environmental factors that modulate plant-pathogen interactions vary in space (Benitez et al. 2013). Indeed, Augspurger & Wilkinson (2007) provide support for species of *Pythium* causing patterns of conspecific negative density dependence in tropical tree species through differential impacts on host species, in concordance with our results (Figure 6). Species of *Phytopythium* may function in a similar manner, but additional pathogenicity experiments are needed to determine the long-term impacts of root infections on seedling survival and growth. These infections could certainly limit the seedlings access to water, nutrients and ability of mycorrhizal fungi to establish.

The diversity of oomycete species was less than observed in similar studies on fungal communities (Sarmiento et al. 2017, Schappe et al. 2020, Spear and Broders 2021). However, the 24 species identified with culture-based baiting and 499 ASVs assigned using metabarcoding is greater than 14 ASVs identified by Oliverio et al. (2020) on BCI, and the 17 OTUs observed by Mahe et al. (2017) on BCI, but similar to the 476 oomycete ASVs found to be associated with adults and seedlings of four tree species on the Gigante Peninsula (Delavaux et al. 2023). The Oliverio et al. (2020) and Mahe et al. (2017) studies used the 28S rRNA gene for designating sequence variants, and this marker is highly conserved among oomycetes and may have under-estimated actual oomycete diversity, whereas this study and the one by Delavaux et al. (2023) used the ITS region with oomycete specific primers (Riit et al. 2016). The use of the ITS region in future metabarcoding experiments is recommended as it is better able to discriminate between species of oomycetes. Further baiting experiments should also be used to recover novel oomycete species and better understand the true genetic diversity within this group of organisms. The need for additional surveys is particularly true for members of *Phytopythium*. The widespread distribution and diversity of undescribed and novel species in the *Phytopythium vexans* complex (PVC) combined with their relative abundance in tropical and subtropical regions demonstrates the need for a better understanding of the ecology of this group of oomycetes. The relative lack of studies on oomycete pathogens in the tropics demonstrates the need for additional sampling and the importance of depositing these strains in a culture collection for future taxonomic investigations.

In summary, the work presented here represents an important step in clarifying the abundance, distribution, and potential role of soil-borne oomycetes in seedling survival and ultimately plant community structure in the lowland tropics. With our field surveys and lab and shadehouse assays, we use: 1) metabarcoding to identify host-specific differential accumulation and composition of soil-borne oomycetes, 2) baiting from soil to quantify disease potential, and 3) pathogenicity on multiple tree species to demonstrate the underappreciated ubiquitous nature of plant pathogenic oomycetes in lowland tropical forests.

## DATA AVAILABILITY STATEMENT

All DNA sequence data generated by this project were deposited in Genbank as accessions OR659743-OR659896 for ITS gene sequences and OR661912-OR662046 for COI gene sequences. Metabarcoding data was deposited in the GenBank SRA database as accession XXXX. A total of 18 oomycete strains (NRRL number in Suppl. Table 1) used in this study were deposited in the ARS Culture Collection and are available to order from https://nrrl.ncaur.usda.gov/.

## ACKNOWLEDGEMENTS

We want to thank Natalie Ferro Lozano for assistance with soil baiting, Joe Sertich for help with collecting soil samples, Tom Usgaard for generating DNA sequence data on Oomycete strains, and Milton Solano for assistance developing the map of BCI with the sampling sites. This work was funded by the United States Department of Agriculture – Agricultural Research Service and with grants from the National Science Foundation (NSF 2039487) and the Simons Foundation (429440).

## AUTHOR CONTRIBUTIONS

K.B., E.R.S and S.J.W conceived the study. K.B., E.R.S., H.E., M.B., M.A.L.P. collected soil, conducted baiting assays, isolated oomycetes from infected tissue. K.B., M.B., M.A.L.P., G.I., extracted DNA from cultures and soil samples, prepared and sequenced DNA and DNA libraries, analyzed ITS and COI sequence data. K.B., H.D.C.B, G.I. analyzed and visualized data. K.B. wrote the first draft. H.D.C.B, E.R.S and S.J.W contributed to subsequent manuscript drafts. All authors have read and approved the final version.

## COMPETING INTERESTS

The authors have declared that no competing interests exist. Mention of trade names or commercial products in this publication is solely for the purpose of providing specific information and does not imply recommendation or endorsement by the U.S. Department of Agriculture. The U.S. Department of Agriculture prohibits discrimination in all its programs and activities on the basis of race, color, national origin, age, disability, and where applicable, sex, marital status, familial status, parental status, religion, sexual orientation, genetic information, political beliefs, reprisal, or because all or part of an individual’s income is derived from any public assistance program. (Not all prohibited bases apply to all programs.) Persons with disabilities who require alternative means for communication of program information (Braille, large print, audiotape, etc.) should contact USDA’s TARGET Center at (202) 720-2600 (voice and TDD). To file a complaint of discrimination, write to USDA, Director, Office of Civil Rights, 1400 Independence Avenue, S.W., Washington, D.C. 20250-9410, or call (800) 795-3272 (voice) or (202) 720-6382 (TDD). USDA is an equal opportunity provider and employer.

## TABLES

**Supplemental Table 1.** Oomycete isolates recovered from soil at 22 sites in a lowland tropical forest in Panama. Isolates were recovered using a leaf baiting method and identified to species based on sequenced data from the internal transcribed spacer (ITS) region of the ribosomal DNA and the cytochrome oxidase I (COI) region of mitochondrial DNA.

**Supplemental Table 2.** List of oomycete strains identified during this study with their corresponding NCBI accession numbers for the internal transcribed spacer (ITS) region of the ribosomal DNA and the cytochrome oxidase I (COI) region of mitochondrial DNA sequence data.

**Supplemental Table 3.** Results of ASV sequences from this study blasted against the ITS sequences from oomycetes recovered from the baiting assay using BLASTn with a 99% match identity and 50% coverage to ensure the accuracy of the matches.

## SUPPLEMENTAL FIGURES

**Supplemental Figure 1.**
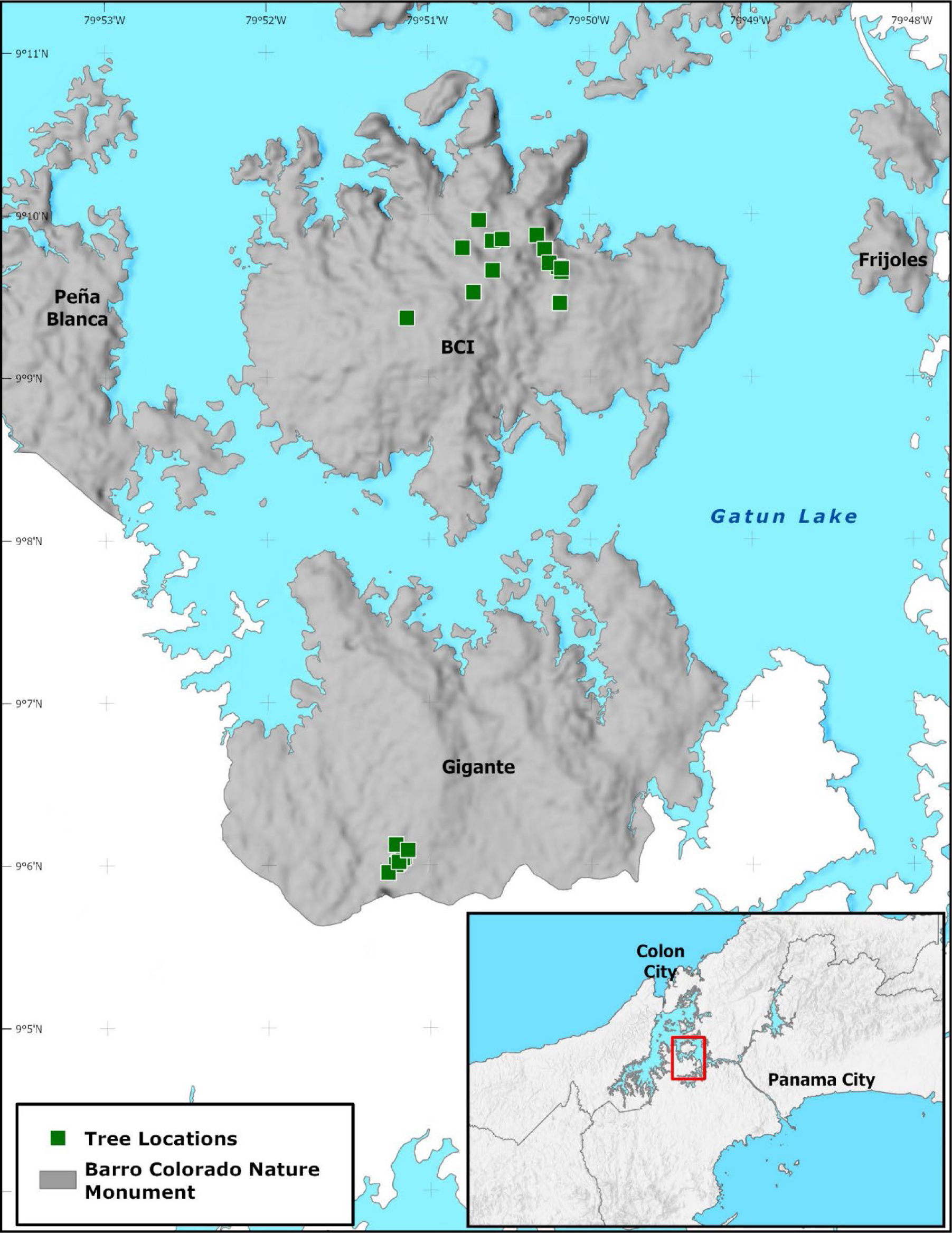
Location of sample sites on Barro Colorado National Monument. Green squares designate individual sampling locations.

**Supplemental figure 2.**
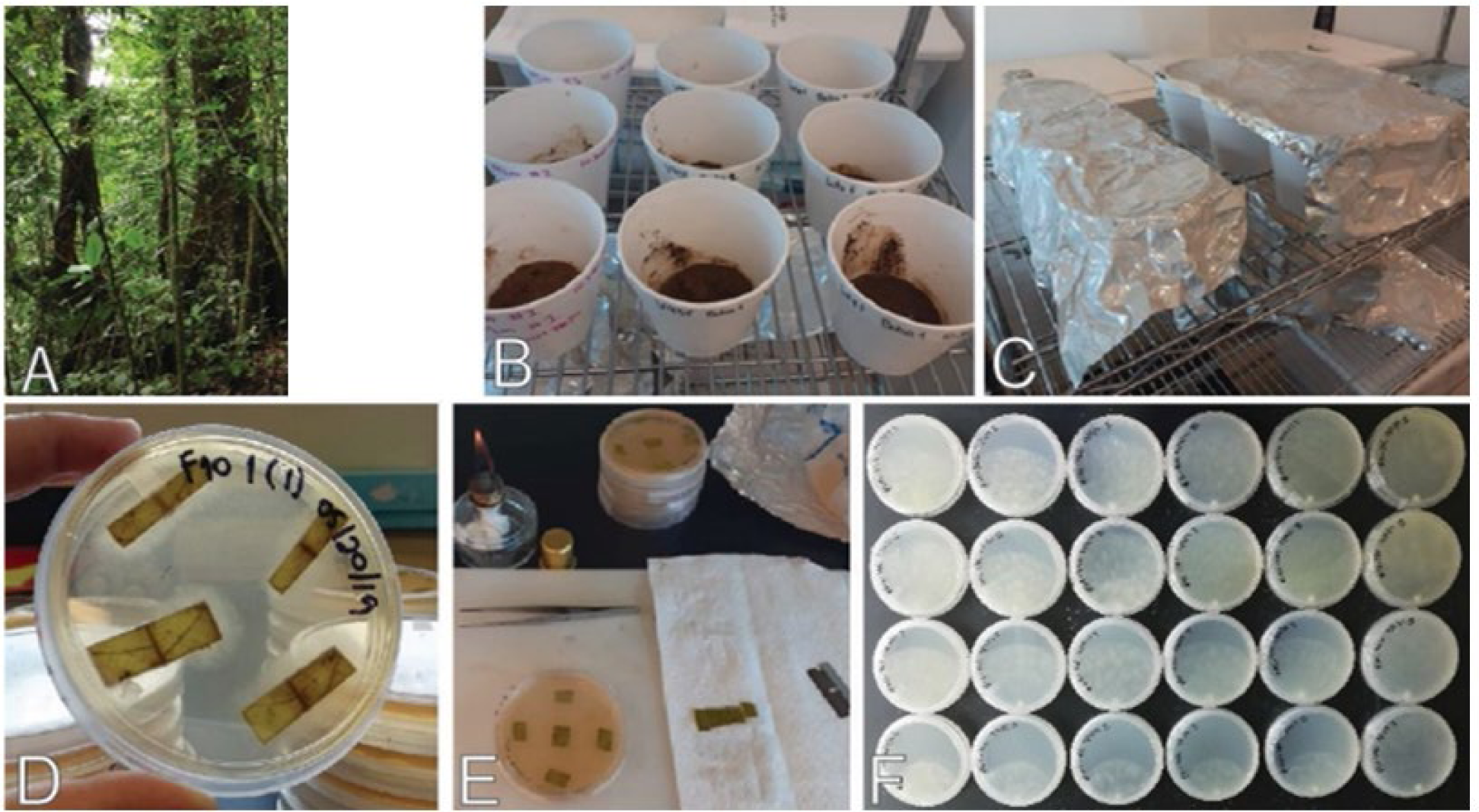
The oomycete baiting process included A) collecting soil from sites on BCI, B) placing 50 g of soil per sample in an opaque container adding 250 ml of sterile water and surface sterilized baits (leaf pieces), C) incubating soil and baits in the dark at room temperature, D) plating baits on antibiotic amended media to allow oomycetes to grow from infected leaves, and F) hyphal-tipping cultures growing from bait leaves to start pure cultures of those isolates.

**Supplemental Figure 3.**
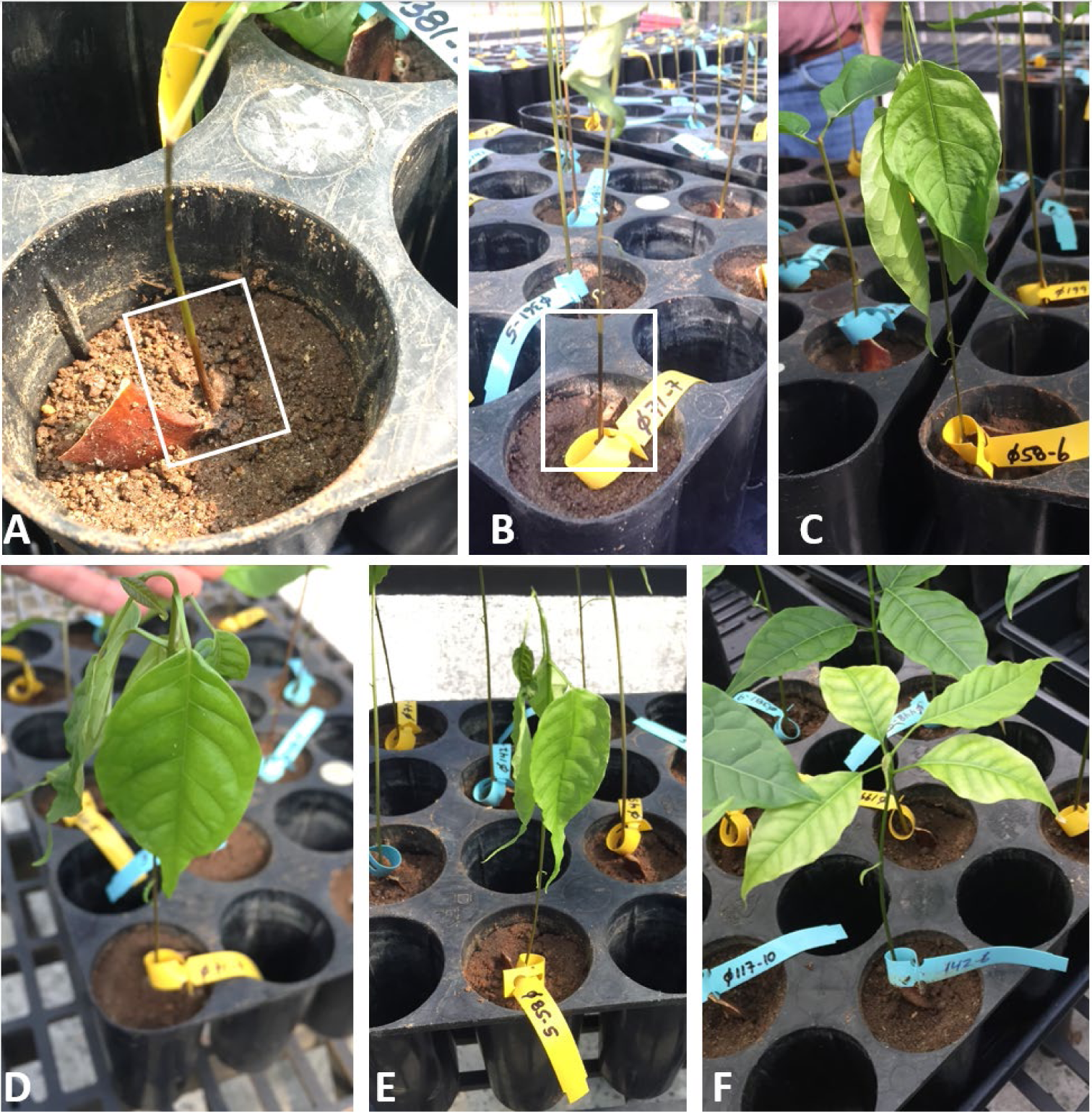
Pathogenicity experiment and symptoms expressed and scored using the ordi including a-b) stem lesions, c-e) wilting and f) chlorosis on *Swietenia macrophylla* seedlings.

**Supplemental Fig 4.**
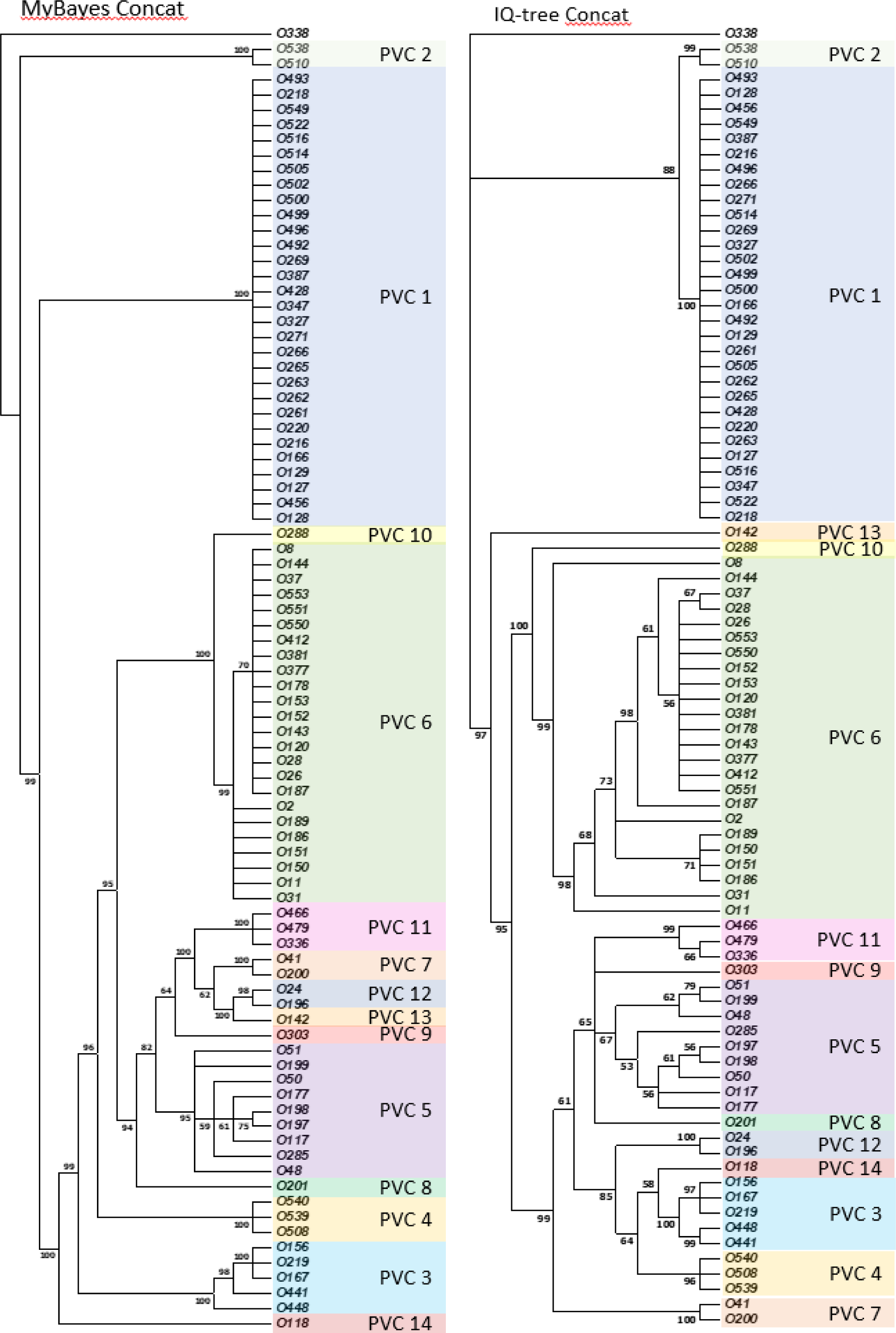
Phylogenetic tree of all isolates sequenced (ITS + CoxI) during this study made using a) Bayesian and b) maximum likelihood algorithms

## Notes

### Competing Interest Statement

The authors have declared no competing interest.

## LITERATURE CITED

Abarenkov, K., R. H. Nilsson, K.-H. Larsson, Andy F. S. Taylor, Tom W. May, T. G. Frøslev, J. Pawlowska, B. Lindahl, K. Põldmaa, C. Truong, D. Vu, T. Hosoya, T. Niskanen, T. Piirmann, F. Ivanov, A. Zirk, M. Peterson, Tanya E. Cheeke, Y. Ishigami, Arnold T. Jansson, Thomas S. Jeppesen, E. Kristiansson, V. Mikryukov, Joseph T. Miller, R. Oono, Francisco J. Ossandon, J. Paupério, I. Saar, D. Schigel, A. Suija, L. Tedersoo, and U. Kõljalg. 2023. The UNITE database for molecular identification and taxonomic communication of fungi and other eukaryotes: sequences, taxa and classifications reconsidered. Nucleic Acids Research.

Agrios, G. A. 2005. Plant Pathology. Academic Press.

Ampt, E. A., J. van Ruijven, M. P. Zwart, J. M. Raaijmakers, A. J. Termorshuizen, and L. Mommer. 2022. Plant neighbours can make or break the disease transmission chain of a fungal root pathogen. New Phytologist 233: 1303–1316.

Augspurger, C. K., and C. K. Kelly. 1984. Pathogen mortality of tropical tree seedlings - Experiment studies of the effects of dispersal distance, seedling density, and light conditions. Oecologia 61: 211–217.

Augspurger, C. K., and H. T. Wilkinson. 2007. Host specificity of pathogenic *Pythium* species: Implications for tree species diversity. Biotropica 39: 702–708.

Avila, J. M., A. Gallardo, B. Ibáñez, and L. Gómez-Aparicio. 2016. *Quercus suber* dieback alters soil respiration and nutrient availability in Mediterranean forests. Journal of Ecology 104: 1441–1452.

Bagchi, R., R. E. Gallery, S. Gripenberg, S. J. Gurr, L. Narayan, C. E. Addis, R. P. Freckleton, and O. T. Lewis. 2014. Pathogens and insect herbivores drive rainforest plant diversity and composition. Nature 506: 85-+.

Balci, Y., S. Balci, J. Eggers, W. L. MacDonald, J. Juzwik, R. P. Long, and K. W. Gottschalk. 2007. *Phytophthora* spp. associated with forest soils in eastern and north-central US oak ecosystems. Plant Disease 91: 705–710.

Balci, Y., R. P. Long, M. Mansfield, D. Balser, and W. L. MacDonald. 2010. Involvement of *Phytophthora* species in white oak (*Quercus alba*) decline in southern Ohio. Forest Pathology 40: 430–442.

Baysal-Gurel, F., P. Liyanapathiranage, M. Panth, F. A. Avin, and T. Simmons. 2021. First report of Phytopythium vexans causing root and crown rot on flowering cherry in Tennessee. Plant Disease 105: 232.

Beakes, G. W., S. L. Glockling, and S. Sekimoto. 2012. The evolutionary phylogeny of the oomycete “fungi”. Protoplasma 249: 3–19.

Benitez, M.-S., M. H. Hersh, R. Vilgalys, and J. S. Clark. 2013. Pathogen regulation of plant diversity via effective specialization. Trends in Ecology & Evolution 28: 705–711.

Bily, D., E. Nikolaeva, T. Olson, and S. Kang. 2022. *Phytophthora* spp. associated with Appalachian oak forests and waterways in Pennsylvania, with *P. abietivora* as a pathogen of five native woody plant species. Plant Disease 106: 1143–1156.

Burdon, J., and G. Chilvers. 1975. Epidemiology of damping-off disease (Pythium irregulare) in relation to density of Lepidium sativum seedlings. Annals of applied Biology 81: 135–143.

Callahan, B. J., P. J. McMurdie, M. J. Rosen, A. W. Han, A. J. A. Johnson, and S. P. Holmes. 2016. DADA2: High-resolution sample inference from Illumina amplicon data. Nature methods 13: 581–583.

Camacho, C., G. Coulouris, V. Avagyan, N. Ma, J. Papadopoulos, K. Bealer, and T. L. Madden. 2009. BLAST+: architecture and applications. BMC Bioinformatics 10: 421.

Campoverde, E. V., G. Sanahuja, and A. J. Palmateer. 2017. A high incidence of *Pythium* and *Phytophthora* diseases related to record-breaking rainfall in south Florida. HortTechnology 27: 78–83.

Chernomor, O., A. Von Haeseler, and B. Q. Minh. 2016. Terrace aware data structure for phylogenomic inference from supermatrices. Systematic biology 65: 997–1008.

Cobb, R. C., J. A. Filipe, R. K. Meentemeyer, C. A. Gilligan, and D. M. Rizzo. 2012. Ecosystem transformation by emerging infectious disease: loss of large tanoak from California forests. Journal of Ecology 100: 712–722.

Davidson, J., S. Rehner, M. Santana, E. Lasso, O. Urena de Chapet, and E. Herre. 2000. First report of Phytophthora heveae and Pythium spp. on tropical tree seedlings in Panama. Plant Disease 84: 706–706.

Davidson, J. M. 2000. Pathogen-mediated maintenance of diversity and fitness consequences of near versus far pollination for tropical trees. University of California-Davis.

Deacon, J. W., and S. P. Donaldson. 1993. Molecular recognition in the homing responses of zoosporic fungi, with special reference to *Pythium* and *Phytophthora*. Mycological Research 97: 1153–1171.

Delavaux, C. S., J. K. Angst, H. Espinosa, M. Brown, D. F. Petticord, J. W. Schroeder, K. Broders, E. A. Herre, J. D. Bever, and T. W. Crowther. 2023. Fungal community dissimilarity predicts plant–soil feedback strength in a lowland tropical forest. Ecology In Press: 10.1002/ecy.4200.

Delavaux, C. S., J. L. Schemanski, G. L. House, A. G. Tipton, B. Sikes, and J. D. Bever. 2021. Root pathogen diversity and composition varies with climate in undisturbed grasslands, but less so in anthropogenically disturbed grasslands. The ISME Journal 15: 304–317.

Dominguez-Begines, J., J. M. Avila, L. V. Garcia, and L. Gomez-Aparicio. 2021. Disentangling the role of oomycete soil pathogens as drivers of plant-soil feedbacks. Ecology 102.

Eck, J. L., S. M. Stump, C. S. Delavaux, S. A. Mangan, and L. S. Comita. 2019. Evidence of within-species specialization by soil microbes and the implications for plant community diversity. Proceedings of the National Academy of Sciences 116: 7371–7376.

Eden, M. A., R. A. Hill, and M. Galpoththage. 2000. An efficient baiting assay for quantification of Phytophthora cinnamomi in soil. Plant Pathology 49: 515–522.

Fawke, S., M. Doumane, and S. Schornack. 2015. Oomycete Interactions with Plants: Infection Strategies and Resistance Principles. Microbiology and Molecular Biology Reviews 79: 263–280.

Fichtner, E., G. Browne, M. Mortaz, L. Ferguson, and C. Blomquist. 2016. First report of root rot caused by Phytopythium helicoides on pistachio rootstock in California. Plant Disease 100: 2337–2337.

Garnas, J. R., M. P. Ayres, A. M. Liebhold, and C. Evans. 2011. Subcontinental impacts of an invasive tree disease on forest structure and dynamics. Journal of Ecology 99: 532–541.

Grover, M., and M. Barkoulas. 2021. *C. elegans* as a new tractable host to study infections by animal pathogenic oomycetes. PLOS Pathogens 17: e1009316.

Hersh, M. H., R. Vilgalys, and J. S. Clark. 2012. Evaluating the impacts of multiple generalist fungal pathogens on temperate tree seedling survival. Ecology 93: 511–520.

Jiang, R. H. Y., I. de Bruijn, B. J. Haas, R. Belmonte, L. Löbach, J. Christie, G. van den Ackerveken, A. Bottin, V. Bulone, S. M. Díaz-Moreno, B. Dumas, L. Fan, E. Gaulin, F. Govers, L. J. Grenville-Briggs, N. R. Horner, J. Z. Levin, M. Mammella, H. J. G. Meijer, P. Morris, C. Nusbaum, S. Oome, A. J. Phillips, D. van Rooyen, E. Rzeszutek, M. Saraiva, C. J. Secombes, M. F. Seidl, B. Snel, J. H. M. Stassen, S. Sykes, S. Tripathy, H. van den Berg, J. C. Vega-Arreguin, S. Wawra, S. K. Young, Q. Zeng, J. Dieguez-Uribeondo, C. Russ, B. M. Tyler, and P. van West. 2013. Distinctive expansion of potential virulence grnes in the genome of the oomycete fish pathogen *Saprolegnia parasitica*. PLOS Genetics 9: e1003272.

Jung, T. 2009. Beech decline in Central Europe driven by the interaction between Phytophthora infections and climatic extremes. Forest Pathology 39: 73–94.

Jung, T., T. T. Chang, J. Bakonyi, D. Seress, A. Pérez-Sierra, X. Yang, C. Hong, B. Scanu, C. H. Fu, K. L. Hsueh, C. Maia, P. Abad-Campos, M. Léon, and M. Horta Jung. 2017. Diversity of Phytophthora species in natural ecosystems of Taiwan and association with disease symptoms. Plant Pathology 66: 194–211.

Jung, T., B. Scanu, C. M. Brasier, J. Webber, I. Milenković, T. Corcobado, M. Tomšovský, M. Pánek, J. Bakonyi, C. Maia, A. Bačová, M. Raco, H. Rees, A. Pérez-Sierra, and M. Horta Jung. 2020. A Survey in Natural Forest Ecosystems of Vietnam Reveals High Diversity of both New and Described Phytophthora Taxa including P. ramorum. Forests 11: 93.

Kalyaanamoorthy, S., B. Q. Minh, T. K. Wong, A. Von Haeseler, and L. S. Jermiin. 2017. ModelFinder: fast model selection for accurate phylogenetic estimates. Nature methods 14: 587–589.

Katoh, K., and D. M. Standley. 2013. MAFFT multiple sequence alignment software version 7: improvements in performance and usability. Molecular biology and evolution 30: 772–780.

Keeling, P. J., and F. Burki. 2019. Progress towards the Tree of Eukaryotes. Current Biology 29: R808–R817.

Khew, K. L., and G. A. Zentmyer. 1973. Chemotactic response of zoospores of 5 species of Phytophthora. Phytopathology 63: 1511–1517.

Latijnhouwers, M., W. Ligterink, V. Vleeshouwers, P. van West, and F. Govers. 2004. A G-alpha subunit controls zoospore motility and virulence in the potato late blight pathogen *Phytophthora infestans*. Molecular Microbiology 51: 925–936.

Lebrija-Trejos, E., A. Hernández, and S. J. Wright. 2023. Effects of moisture and density-dependent interactions on tropical tree diversity. Nature 615: 100–104.

Legeay, J., C. Husson, B. Boudier, E. Louisanna, C. Baraloto, H. Schimann, B. Marcais, and M. Buee. 2020. Surprising low diversity of the plant pathogen *Phytophthora* in Amazonian forests. Environmental Microbiology 22: 5019–5032.

Li, Y., Y. Feng, C. Wu, J. Xue, B. Jiao, B. Li, and T. Dai. 2021. First report of phytopythium litorale causing crown and root rot on Rhododendron pulchrum in china. Plant Disease 105: 4173.

Mahe, F., C. de Vargas, D. Bass, L. Czech, A. Stamatakis, E. Lara, D. Singer, J. Mayor, J. Bunge, S. Sernaker, T. Siemensmeyer, I. Trautmann, S. Romac, C. Berney, A. Kozlov, E. A. D. Mitchell, C. V. W. Seppey, E. Egge, G. Lentendu, R. Wirth, G. Trueba, and M. Dunthorn. 2017. Parasites dominate hyperdiverse soil protist communities in Neotropical rainforests. Nature Ecology & Evolution 1.

Mangan, S. A., S. A. Schnitzer, E. A. Herre, K. M. L. Mack, M. C. Valencia, E. I. Sanchez, and J. D. Bever. 2010. Negative plant-soil feedback predicts tree-species relative abundance in a tropical forest. Nature 466: 752–U710.

Martin, M. 2011. Cutadapt removes adapter sequences from high-throughput sequencing reads. EMBnet. journal 17: 10–12.

Masikane, S., J. Jolliffe, L. Swart, and A. McLeod. 2019. Novel approaches and methods for quantifying Phytophthora cinnamomi in avocado tree roots. FEMS Microbiology Letters 366.

McMurdie, P. J., and S. Holmes. 2013. phyloseq: An R Package for Reproducible Interactive Analysis and Graphics of Microbiome Census Data. Plos One 8: e61217.

Nguyen, L.-T., H. A. Schmidt, A. Von Haeseler, and B. Q. Minh. 2015. IQ-TREE: a fast and effective stochastic algorithm for estimating maximum-likelihood phylogenies. Molecular biology and evolution 32: 268–274.

Noireung, P., P. Intaparn, R. Maumoon, T. Wongwan, and C. To–anun. 2020. First Record of Phytopythium vexans causing root rot on Mandarin (Citrus reticulate L. cv. Sainampueng) in Thailand. Plant Pathology & Quarantine 10: 85–90.

Oliverio, A. M., S. Geisen, M. Delgado-Baquerizo, F. T. Maestre, B. L. Turner, and N. Fierer. 2020. The global-scale distributions of soil protists and their contributions to belowground systems. Science Advances 6.

Packer, A., and K. Clay. 2000. Soil pathogens and spatial patterns of seedling mortality in a temperate tree. Nature 404: 278–281.

Packer, A., and K. Clay. 2004. Development of negative feedback during successive growth cycles of black cherry. Proceedings of the Royal Society B-Biological Sciences 271: 317–324.

Riit, T., L. Tedersoo, R. Drenkhan, E. Runno-Paurson, H. Kokko, and S. Anslan. 2016. Oomycete-specific ITS primers for identification and metabarcoding. MycoKeys 14: 17–30.

Robideau, G. P., A. de Cock, M. D. Coffey, H. Voglmayr, H. Brouwer, K. Bala, D. W. Chitty, N. Desaulniers, Q. A. Eggertson, C. M. M. Gachon, C. H. Hu, F. C. Kupper, T. L. Rintoul, E. Sarhan, E. C. P. Verstappen, Y. H. Zhang, P. J. M. Bonants, J. B. Ristaino, and C. A. Levesque. 2011. DNA barcoding of oomycetes with cytochrome c oxidase subunit I and internal transcribed spacer. Molecular Ecology Resources 11: 1002–1011.

Rollins, L., K. Coats, M. Elliott, and G. Chastagner. 2016. Comparison of Five Detection and Quantification Methods for *Phytophthora ramorum* in Stream and Irrigation Water. Plant Disease 100: 1202–1211.

Ronquist, F., M. Teslenko, P. Van Der Mark, D. L. Ayres, A. Darling, S. Höhna, B. Larget, L. Liu, M. A. Suchard, and J. P. Huelsenbeck. 2012. MrBayes 3.2: efficient Bayesian phylogenetic inference and model choice across a large model space. Systematic biology 61: 539–542.

Ruffner, B., D. Rigling, and C. N. Schoebel. 2019. Multispecies *Phytophthora* disease patterns in declining beech stands. Forest Pathology 49: e12514.

Sarker, S. R., J. McComb, G. E. S. J. Hardy, and T. I. Burgess. 2023. Sample volume affects the number of *Phytophthora* and *Phytopythium* species detected by soil baiting. European Journal of Plant Pathology 166: 303–313.

Sarmiento, C., P.-C. Zalamea, J. W. Dalling, A. S. Davis, S. M. Stump, J. M. U’Ren, and A. E. Arnold. 2017. Soilborne fungi have host affinity and host-specific effects on seed germination and survival in a lowland tropical forest. Proceedings of the National Academy of Sciences of the United States of America 114: 11458–11463.

Sato, T., T. Egusa, K. Fukushima, T. Oda, N. Ohte, N. Tokuchi, K. Watanabe, M. Kanaiwa, I. Murakami, and K. D. Lafferty. 2012. Nematomorph parasites indirectly alter the food web and ecosystem function of streams through behavioural manipulation of their cricket hosts. Ecology Letters 15: 786–793.

Schappe, T., F. E. Albornoz, B. L. Turner, and F. A. Jones. 2020. Co-occurring fungal functional groups respond differently to tree neighborhoods and soil properties across three tropical rainforests in Panama. Microb Ecol 79: 675–685.

Sedio, B. E., and A. M. Ostling. 2013. How specialised must natural enemies be to facilitate coexistence among plants? Ecology Letters 16: 995–1003.

Simon, T. E., R. Le Cointe, P. Delarue, S. Morlière, F. Montfort, M. R. Hervé, and S. Poggi. 2014. Interplay between Parasitism and Host Ontogenic Resistance in the Epidemiology of the Soil-Borne Plant Pathogen Rhizoctonia solani. PLOS ONE 9: e105159.

Spear, E. R., and K. D. Broders. 2021. Host-generalist fungal pathogens of seedlings may maintain forest diversity via host-specific impacts and differential susceptibility among tree species. New Phytologist 231: 460–474.

Spear, E. R., P. D. Coley, and T. A. Kursar. 2015. Do pathogens limit the distributions of tropical trees across a rainfall gradient? Journal of Ecology 103: 165–174.

Spies, C. F. J., A. M. Grooters, C. A. Lévesque, T. L. Rintoul, S. A. Redhead, S. L. Glockling, C.-y. Chen, and A. W. A. M. de Cock. 2016. Molecular phylogeny and taxonomy of Lagenidium-like oomycetes pathogenic to mammals. Fungal Biology 120: 931–947.

Sturrock, R., S. Frankel, A. Brown, P. Hennon, J. Kliejunas, K. Lewis, J. Worrall, and A. Woods. 2011. Climate change and forest diseases. Plant Pathology 60: 133–149.

Tamura, K., G. Stecher, and S. Kumar. 2021. MEGA11: molecular evolutionary genetics analysis version 11. Molecular biology and evolution 38: 3022–3027.

Thompson, S., S. Levin, and I. Rodriguez-Iturbe. 2013. Linking plant disease risk and precipitation drivers: a dynamical systems framework. The American Naturalist 181: E1–E16.

Wang, K., Y. Xie, G. Yuan, Q. Li, and W. Lin. 2015. First report of root and collar rot caused by Phytopythium helicoides on kiwifruit (Actinidia chinensis). Plant Disease 99: 725.

Wang, Q., G. M. Garrity, J. M. Tiedje, and J. R. Cole. 2007. Naïve Bayesian Classifier for Rapid Assignment of rRNA Sequences into the New Bacterial Taxonomy. Applied and Environmental Microbiology 73: 5261–5267.

White, T. J., T. Bruns, S. Lee, and J. Taylor. 1990. Amplification and direct sequencing of fungal ribosomal RNA genes for phylogenetics. PCR protocols: a guide to methods and applications 18: 315–322.

Windsor, D. M. 1990. Climate and moisture variability in a tropical forest: long-term records from Barro Colorado Island, Panama. Smithsonian contributions to the earth sciences.

Yavitt, J. B. 2024. Soils of Barro Colorado Island. Pages XXX-XXX *in* H. C. Muller-Landau and S. J. Wright, editors. The First 100 Years of Research on Barro Colorado Island: Plant and Ecosystem Science. Smithsonian Institution Scholarly Press, Washington D.C.

